# Electrical stimulation directs formation of perfused vasculature in engineered tissues

**DOI:** 10.1101/2025.08.28.672965

**Authors:** Katarzyna A. Grzelak, Ashley D. Westerfield, Vardhman Kumar, Kasturi Chakraborty, Navaneeth Krishna Rajeeva Pandian, Christopher S. Chen, Sangeeta N. Bhatia

## Abstract

Effective, rapid and functionally perfusable vascularization remains a major challenge in tissue engineering. Current approaches to generate vasculature *in vitro* require multipart fabrication methods or complex and costly media supplements, limiting their scalability. Here, we demonstrate that exogenous electrical stimulation (estim) offers a promising alternative by enhancing 3D vasculogenesis in engineered human tissues. Exposing 3D endothelial-fibroblast cocultures to pulsed estim promoted the formation of dense and branched vascular networks. In a microfluidic device model, we show that estim induces the formation of an interconnected vascular network that can be perfused, whereas unstimulated control networks remained less mature. Importantly, we demonstrate that upon implantation, estim-pretreated vascular grafts exhibit elevated anastomosis with host and perfusion with blood relative to the untreated grafts. In addition, we use estim to promote engraftment of a vascularized 3D liver construct. Mechanistically, we find that estim induces membrane hyperpolarization in endothelial cells via voltage-gated potassium (K_V_) channels. Inhibiting K_V_ channels abrogated estim’s pro-vasculogenic effects in endothelial cells. Conversely, pharmacologically activating hyperpolarization induced endothelial responses even without estim, directly linking K_V_ channel-mediated hyperpolarization as a key mechanism by which estim drives vascular assembly and function. Ultimately, our work establishes estim as a new orthogonal approach to promote formation of perfusable vasculature in engineered tissues.

## Introduction

Tissue-engineered constructs rely on vasculature to mediate transport of oxygen and nutrients critical for tissue viability. This requirement for vascularized tissue models and grafts is particularly relevant for metabolically demanding tissues like liver or heart, which contain up to 3000 vessels/mm^3^[1]. Current vascular engineering approaches, ranging from top-down fabrication methods like 3D printing and sacrificial molding to bottom-up strategies involving endothelial cell self-assembly into vessels using co-culture and pro-angiogenic cues have enabled progress in forming microvascular structures (reviewed in [2], [3]). While significant advancements have been made in promoting vascular assembly, maturation, and remodeling, current techniques often face limitations, and achieving fully perfusable networks of constructs *in vitro* and their effective integration with host vasculature upon implantation remains a challenge. Therefore, developing complementary or alternative strategies that enable the formation of stable, functional, and host-connected vasculature in engineered tissues would enable advancement of the field of tissue engineering.

Bioelectric signaling plays a pivotal role in several physiological processes, including tissue regeneration, cellular migration, and organ development [4], [5]. As a result, the application of exogenous electrical stimulation to clinical settings has been suggested as a means of enhancing wound healing and regeneration. Several studies report improvements in vascularization and nascent vessel formation following *in vivo* electrical stimulation of various tissues [6], [7], [8], [9]. This suggests that electrical stimulation may be a promising, yet underexplored approach in promoting perfusion and vascular integration in the context of engineered tissue. Studies of electrical stimulation on endothelial cells in two-dimensional (2D) and 2.5-dimensional (2.5D) cultures where cells are cultured on Matrigel have shown that estim can enable early vasculogenesis [10], [11], [12], [13], [14], supporting the notion that electrical stimulation can directly influence endothelial cells However, the complex process of assembling endothelial cells into functional, perfusable vasculature has not yet been addressed. Mechanistically, it has been reported in 2D studies that electrical stimulation increases directional galvanotaxis[15], proliferation and results in pro-angiogenic endothelial phenotype through increased production of vascular endothelial growth factor A (VEGFA), activation of vascular endothelial growth factor receptors 1 and 2 (VEGFR1 and VEGFR2), C-X-C chemokine receptors (CXCRs), phosphoinositide 3-kinase/protein kinase B (PI3K/Akt) and mitogen-activated protein kinase/ extracellular signal-regulated kinase (MAPK/ERK) signaling[10], [13], [16], [17], [18].While these studies have begun to explore downstream molecular pathways activated by electrical stimulation, the upstream biophysical effects, such as changes of membrane potential (V_mem_), remain unclear. As a key orchestrator of numerous cellular processes, V_mem_ may serve as an overarching physical regulator linking electrical cues to diverse molecular responses.

In this study, we employ direct current (DC)-pulsed electrical stimulation (estim) to promote a pro-angiogenic state in human umbilical vein endothelial cells (HUVECs). Using this method, we engineer vasculature within *in vitro* tissue constructs, and demonstrate that these electrically stimulated networks are not only structurally enhanced, i.e. denser and more branched, but also functionally perfusable. Furthermore, in an *in vivo* implantation model, we show that the estim-pretreated human vasculature anastomoses with the mouse host to a greater extent than non-treated control within vascular as well as hepatocyte-laden constructs. Mechanistically, we demonstrate that V_mem_ increases with estim, specifically via increased transport through voltage-gated potassium (K_V_) channels, which in turn contributes to the observed functional improvements in vascular network assembly. This report extends prior work by showing estim-mediated enhancement of vasculogenic responses and perfusion in a tissue-engineered 3D construct. Additionally, our study establishes the use of estim to generate functional 3D vasculature via control over membrane hyperpolarization induced by K_V_ channels.

## Results

### Electrical stimulation promotes vasculogenic responses and improves assembly of initial vascular plexus

In order to study how electrical stimulation (estim) affects vascular endothelium, we built a device that consists of gold-plated electrodes positioned at opposite ends of the chamber, connected to a waveform generator delivering electric pulses (**Figure 1A**). We designed our device to be modular and therefore easily amenable to various culture conditions, allowing for controlled application of electric fields (EFs) to a variety of 2D- and 3D-cultured cells.

**Figure 1.**
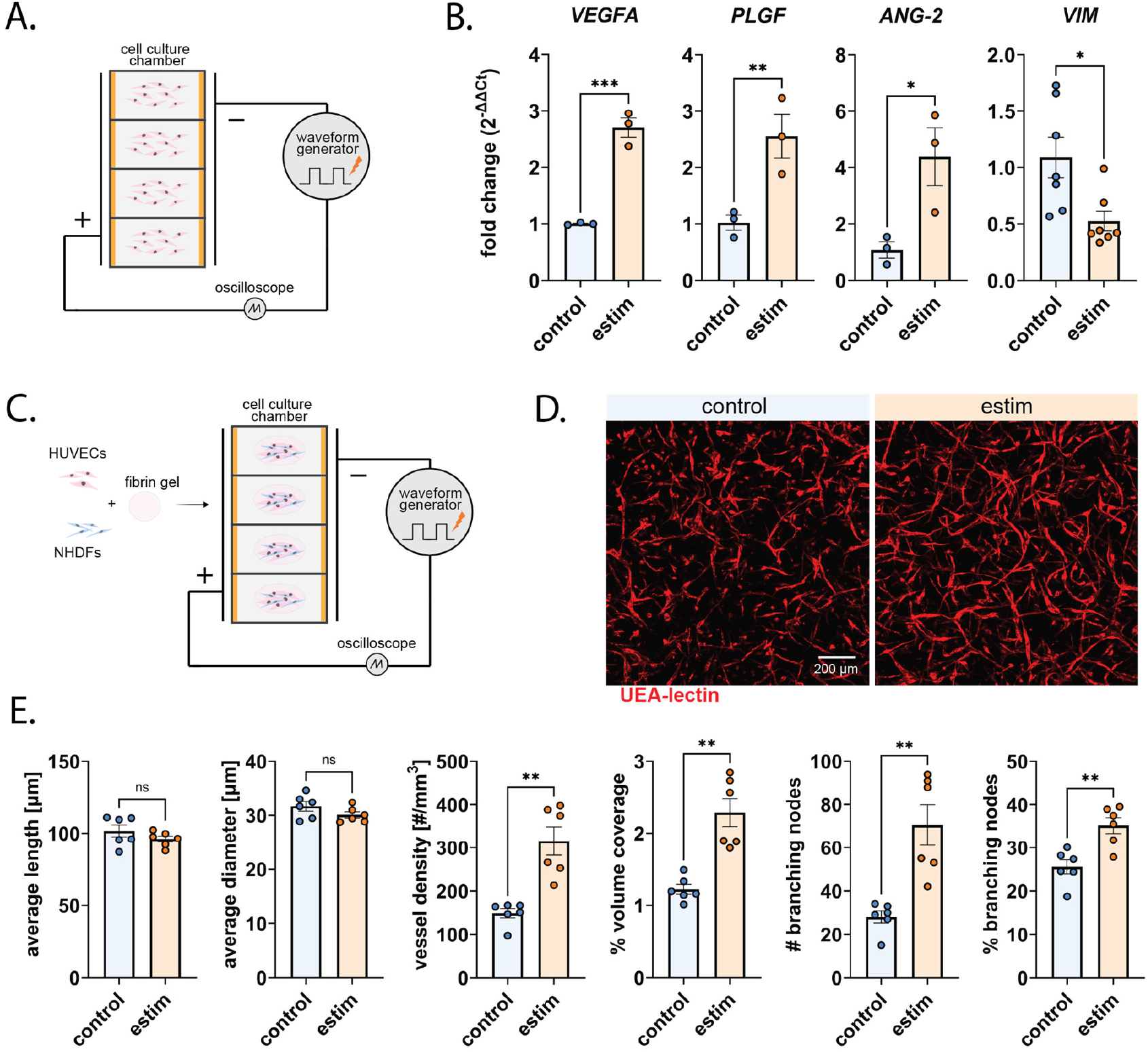
Electrical Stimulation Enhances Vascular Network Formation in 2D and 3D Cultures. a) Schematic of the electrical stimulation (estim) device, showing the electrodes positioned at opposite ends of the chamber and connected to a waveform generator with cells plated in each individual chamber; b) qPCR analysis of gene expression in 2D HUVEC cultures subjected to estim, at least three wells per condition; c) schematic of the seeding workflow for 3D cultures in estim device; d) representative maximum intensity projections of confocal images of 3D HUVEC-fibroblast co-cultures in fibrin hydrogel, stained with UEA I-lectin to visualize the endothelial network; e) Quantification of 3D network properties, six hydrogel fields of view per condition; unpaired t-test, ns not significant, * p<0.05. ** p<0.01, *** p<0.001.

We first optimized the pulse parameters to ensure that the estim did not compromise cell viability, while staying within a physiologically relevant range [5], [19]. In a 2D monolayer culture of HUVECs, the maximum tolerated pulse parameters delivered by our system were found to be 0.2 ms pulse width, 1 Hz frequency, and 2.5 V/cm electric field strength. These estim conditions were selected based on viability assessment, which showed neither significant cell death at these settings (**Figure S1**), nor detectable electrolysis byproducts that would change the pH or temperature of the media (**Figure S1**).

Using optimized parameters, we explored the effects of estim on the expression of pro-angiogenic growth factors, given that VEGFA has been reported to be upregulated under electrical stimulation[10], [13], [20]. We observed that upon receiving estim, gene expression of *VEGFA*, angiopoietin-2 (*Ang-2*), and placental growth factor (*PLGF*) all increased, suggesting a shift towards a pro-vasculogenic profile in the endothelial cells (**Figure 1B**). Notably, we also saw a reduction in vimentin (*VIM*) expression, a mesenchymal marker, further indicating that estim promotes endothelial phenotype in HUVECs.

Next, we moved to a 3D culture system using fibrin hydrogels to begin investigating the effects of estim on the initial steps of vascular network formation. We chose fibrin as the support matrix due to its inherent pro-angiogenic properties and biodegradability, allowing for vascular remodeling[21]. We opted to include human fibroblasts in the system as a proxy for vascular mural cells which were previously shown to support vascular network formation and stability[22], [23]. We therefore co-cultured HUVECs and neonatal human dermal fibroblasts (NHDFs) encapsulated in fibrin domes for 4 days in the presence or absence of estim (**Figure 1C**). At the end of the estim, the samples were fixed and UEA I-lectin staining was used to visualize the human endothelial cells (**Figure 1D**) and assess the extent of vascular network formation. The resulting network characteristics were quantified using an unbiased image analysis script, by measuring average length and diameter of the links, defined as the vessels between nodes, vessel density, volume coverage, and number and percentage of branching nodes. Estim significantly enhanced vessel density and volume coverage, as well as the number of branching nodes compared to controls (**Figure 1E**). However, we did not observe any significant changes in the average length or diameter of the individual vessels. These findings suggest that electrical stimulation promotes the formation of denser, and more interconnected vascular networks without altering the basic structural characteristics of the links.

### Electrical stimulation enhances perfusion of 3D self-assembled vessels in a microfluidic device

Having demonstrated that estim promotes the formation of denser vascular networks, we next sought to investigate the functionality of these vessels. To assess this, we adapted our estim system to a microfluidic device designed to study vasculogenesis *in vitro* by incorporating electrodes (**Figure 2A**). The electrodes were inserted into their respective slots, separated by the central compartment of the device, which was seeded with HUVEC-NHDF co-cultures in fibrin. This compartment is where vascular assembly can be monitored. The generated vasculature progressively connects to the two top-down fabricated channels, 250 µm in diameter, which connect to media ports that provide a means of perfusion of the device (**Figure 2A**). This design allowed for precise control over the culture environment, enabling us to evaluate the effect of estim treatment on the formation of functional, perfusable vascular networks.

**Figure 2.**
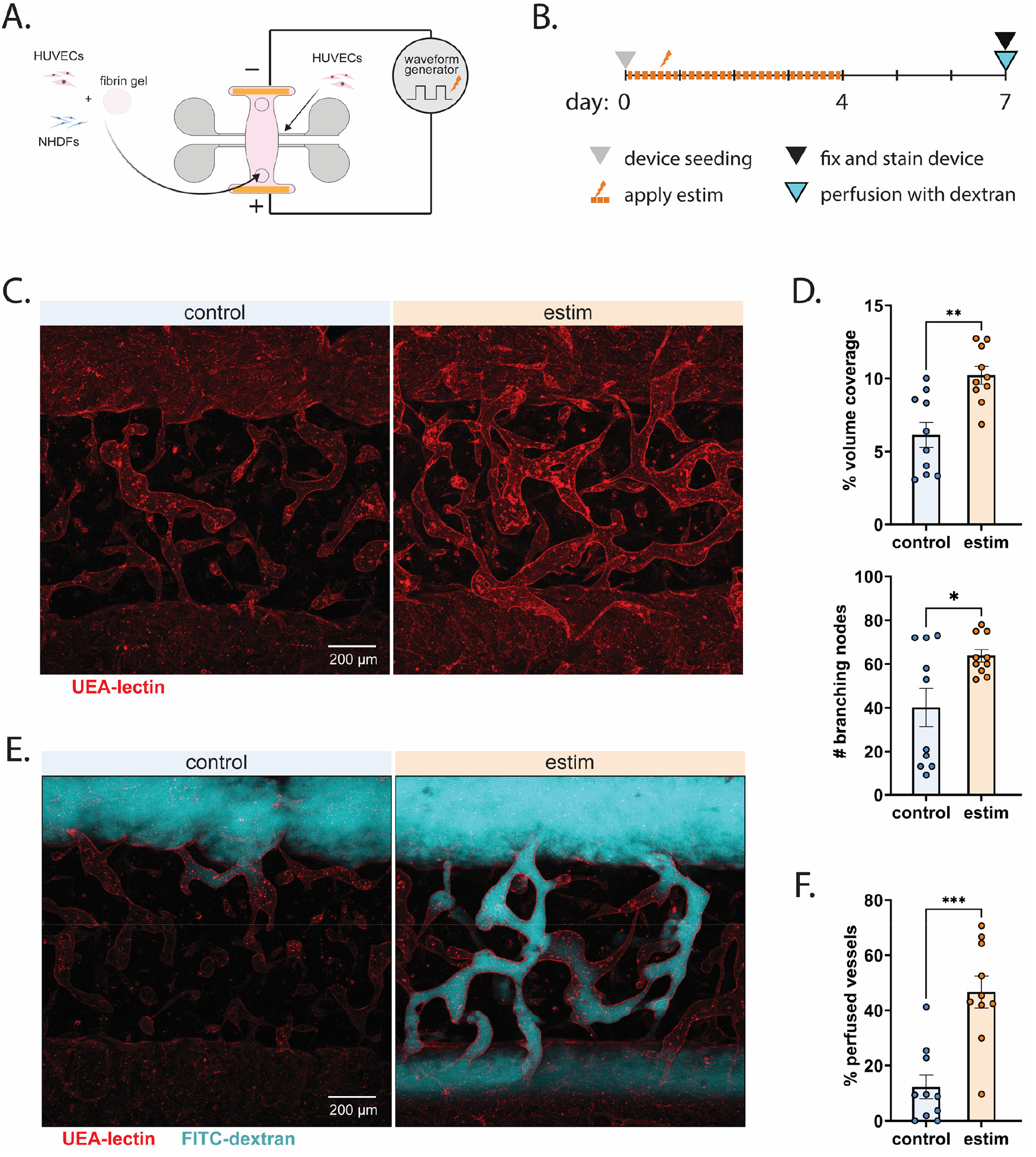
Estim Induces Functional Vascularization in a 3D Microfluidic Vascularization Model. a) Schematic representation of the microfluidic device used for studying vasculogenesis in vitro under electrical stimulation; b) experimental timeline; c) representative max projection images of microfluidic devices at day 7, showing vascular network formation in HUVEC-NHDF co-cultures stained with UEA I-lectin; d) quantification of vascular network properties in the microfluidic devices, branching node count, and network volume coverage; e) representative max projection images showing dextran perfusion in microfluidic devices; f) quantification of dextran perfusion in the microfluidic devices, showing perfused vessels in estim-treated cultures; unpaired t-test, replicates indicate ten hydrogel regions, * p<0.05. ** p<0.01, *** p<0.001.

HUVEC-NHDF co-cultures were seeded in the device (**Figure 2A**), and initially cultured in estim for four days, followed by a recovery period without estim for three days (**Figure 2B**). After a week of culture, we observed an improvement in vessel density within the estim devices (**Figure 2C**), consistent with our finding in 3D fibrin domes. Quantification of confocal images confirmed that HUVECs in estim-treated devices formed more interconnected and dense vascular networks compared to controls, as indicated by an increase in volume coverage and number of branching nodes (**Figure 2D**). To confirm that the networks formed under estim were functional, we analyzed dextran perfusion through the vascular networks (**Figure 2E**), where we saw continuous connection between the newly formed vessels and the top-down fabricated channels in the estim group.

Quantification of the images revealed a significant improvement in vessel perfusion in the estim-treated cultures (**Figure 2F**). The data show that estim promotes the formation of highly perfused vessels, further supporting the notion that estim enhances vascular functionality *in vitro* in addition to increasing the density and complexity of the formed vessels.

The increase in vessel density and perfusion may be due to increased production of factors such as *VEGFA* and *ANG-2* (**Figure 1B**). While these factors are reported to promote vasculogenesis, they are also known to contribute to vascular leakiness[24]. To address this, we examined vessel permeability using low molecular weight fluorescent dextran (70 kDa) in the top-down fabricated vessels. Despite the observed improvements in network density and perfusion, no significant differences in vessel permeability were detected between estim-treated and control groups (**Figure S2**). Thus, estim promotes vascular development and perfusion, without affecting vessel permeability.

### Electrical stimulation facilitates anastomosis and perfusion by mouse systemic circulation of implanted engineered human tissues

Since estim significantly improved the density, interconnectivity, and perfusability of generated vessels in an *in vitro* vasculogenesis model, we sought to adapt our system to an *in vivo* context as a simple and easily adaptable tool to vascularize engineered tissues and improve their engraftment. We generated HUVEC-NHDF fibrin grafts and conditioned them in estim for two days *in vitro* before implanting them subcutaneously into Balb/c nude mice for two weeks (**Figure 3A**). After 14 days, the host mice were injected with lysine-fixable fluoresent dextran intravenously to allow for visualization of anastomosed and lumenized vasculature. The explanted grafts were stained for human endothelial cells (**Figure 3B**), and unbiased image analysis revealed an improvement in volume coverage and branching of the networks in the estim pre-treated grafts compared to control (**Figure 3C**). This result was notable for indicating prolonged benefits of estim after only a short duration of estim pre-treatment with the benefits of two days of estim lingering for at least two subsequent weeks. Interestingly, FITC-dextran perfusion showed that at the time of explantation, only one of the control grafts had detectable perfused vessels, while three out of four explanted pre-estimed grafts were majorly perfused (**Figure 3D**). Image analysis revealed a significant increase in perfusion of the estim pre-treated grafts (**Figure 3E**). Previous reports have shown that vascular grafts generated with 1 day *in vitro* pre-culture, while initially anastomose following implantation, they subsequently accumuate clots and remain unperfusable whereas mature networks precultured for 14 days persist *in vivo* [25]. However, estim appears to promote more persistent and stable vessel perfusion after as little as 2 days of pretreatment.

**Figure 3.**
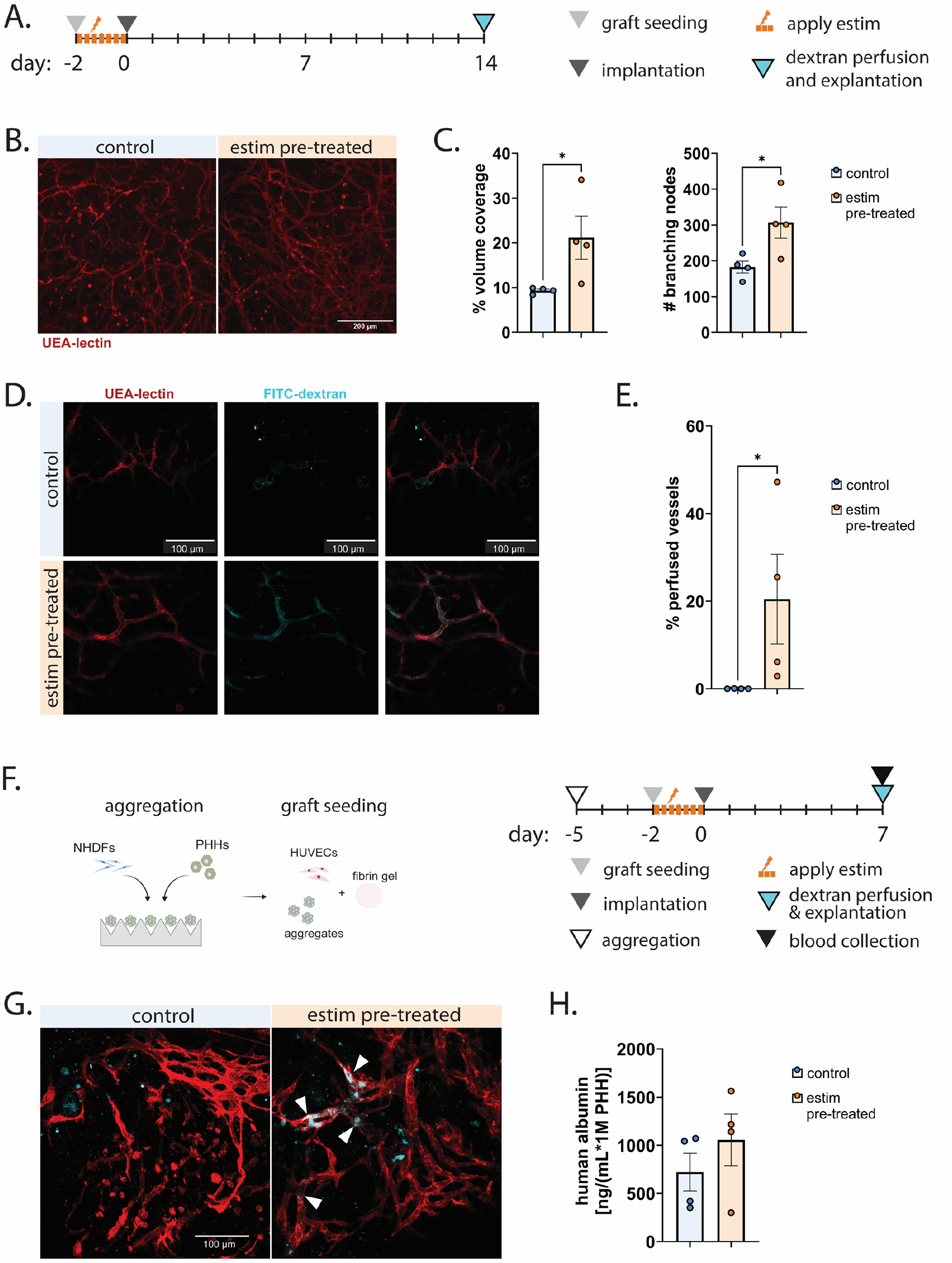
Estim-Fabricated Vessels Improve Graft Vascularization and Perfusion Upon Implantation. a) schematic of experimental workflow: vascular-only graft seeding and subcutaneous implantation; b) representative maximum intensity projections of confocal images of grafts explanted at day 14 following subcutaneous implantation; c) quantification of graft human vascuature properties: volume coverage and branching; d) maximum intensity projections of concofal images of fluorescent dextran within the anastomosed human vasculature, perfusable at day 14 following implantation; e) quantification of %perfused human vasculature in confocal images taken of explant grafts; f) schematic of experimental workflow: generation of hepatic grafts with estim pre-treated vasculature, and intraperitoneal implantation prior to explantation at day 7; g) maximum intensity projections of confocal images of dextran-perfused human vasculature in the hepatic grafts, white arrows point to perfused human vasculature; h) human albumin ELISA results from the implanted hepatic grafts. For c) and e) replicates reported as average of two to three fields of view per each graft (four total per condition), unpaired t-test, * p<0.05; for h) replicates reported as averaged values per mouse with graft (four per condition).

Given the promise of estim in improving engineered tissue vascularization, we assessed our system’s ability to vascularize and improve engraftment of a highly metabolically active parenchyma. To do this, we focued on engineered hepatic tissue. We generated a liver graft by encapsulating spheroids of primary human hepatocytes (PHH) and NHDFs in fibrin hydrogel together with HUVECs and NHDFs. The grafts were pre-treated with estim prior to implantation into the mouse mesenteric fatpad. Notably, estim exposure of PHH within the *in vitro* constructs did not negatively impact the functionality of these cells (**Figure S3**). Following a dextran perfusion, the grafts were explanted after 7 days. Actively perfused vessels were only detected in estim pre-treated grafts (**Figure 3G**) indicating a more anastomosed and lumenized vasculature that can be attributed to the preconditioning with estim. Moreover, when probing for the presence of human albumin in the mouse plasma, we detected an upward, albeit non-significant trend in the estim-pretreated compatred to control grafts at the 7 days timepoint (**Figure 3H**). It is possible that at later timepoints we would see further increase in albumin production in the estim-vascularized condition. Therefore, estim treatment confers perfusion benefit in hepatic grafts without causing adverse effects on hepatocytes.

### Electrical stimulation leads to membrane hyperpolarization

Having shown that estim leads to enhanced vascular formation both *in vitro* and *in vivo*, as well as promotes increased host-graft anastomosis, we aimed to elucidate the mechanisms that enable electrical signals to be transduced into the functional outcomes in conventionally non-electroactive cells.

While a number of mechanisms have been previously explored [26], [27], these studies focused on intracellular cascades and did not directly connect them with electrophysiological processes at play. Membrane potential itself has been suggested as an instructive cue for the cell [28], [29]. We therefore hypothesized that membrane polarization changes upon treatment with estim and may be upstream of the biological signaling cascades studied previously.

To test this hypothesis, we investigated how V_mem_ changes upon estim treatment. HUVECs stained with bis-(1,3-dibutylbarbituric acid)trimethine oxonol (DiBAC_4_(3)), a membrane potential dye, revealed a loss of signal under estim (**Figure 4A**), indicating hyperpolarization of the cell membrane. Interestingly, NHDFs present in the 3D models of **Figure 1** and **2**, did not change their V_mem_ when exposed to estim (**Figure S4**). Quantification of the HUVEC DiBAC_4_(3) fluorescence (**Figure 4B**) confirmed a significant decrease in fluorescence over the course of 2 hours of estim treatment, consistent with the pattern of membrane hyperpolarization.

**Figure 4.**
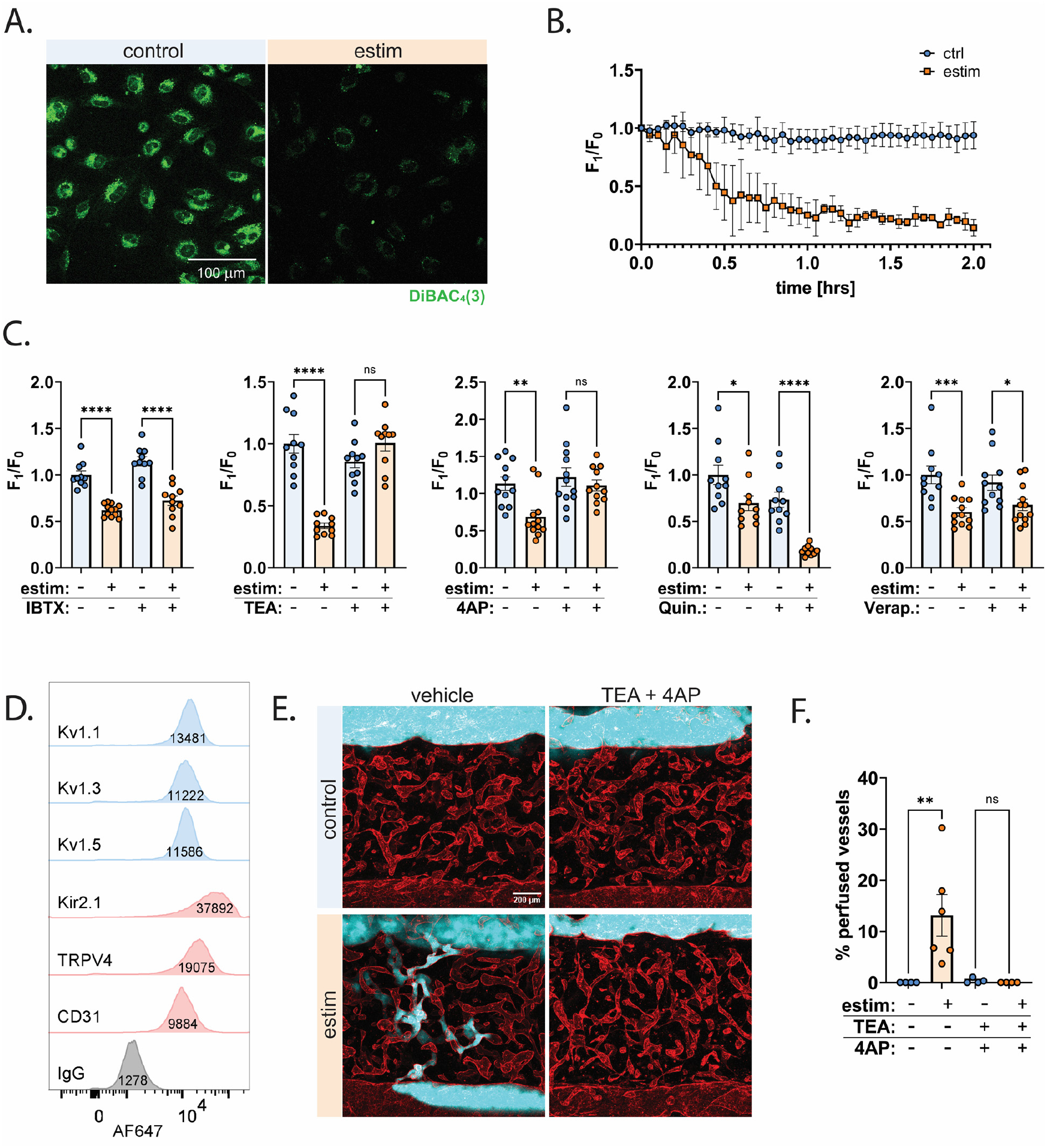
Voltage-Gated Potassium Channels Mediate Estim-Induced Hyperpolarization and Vascularization. a) Representative images of HUVECs stained with DiBAC_4_(3); b) quantification of DiBAC_4_(3) fluorescence changes in HUVECs subjected to electrical stimulation over the course of 2 hours at 3 min intervals, note the decrease in fluorescence indicating membrane hyperpolarization, averaged fluorescence values from three fields of view; c) quantification of DiBAC4(3) fluorescence in ten individual cells at 2 h in 2D HUVEC cultures treated with electrical stimulation and selected ion channel inhibitors (IBTX – iberiotoxin, TEA - tetraethylammonium, 4-AP – 4-aminopyridine, Quin. - quinidine, Ver. - verapamil); d) comparison of fluorescence intensity of antibodies staining for voltage-gated potassium channels (Kv 1.1, 1.3, 1.5), Kir2.1, TRPV4, and CD31 compared to IgG control, obtained via flow cytometry; e) representative images of microfluidic devices showing the effect of potassium channel blockers (TEA and 4-AP) on vascular architecture in the presence of electrical stimulation. f) quantification of dextran perfusion in microfluidic devices treated with electrical stimulation and potassium channel blockers in at least four hydrogel regions per condition. One-way ANOVA, ns - not significant, * p< 0.05, ** p<0.01, *** p<0.001, ****p<0.0001.

To further investigate whether this hyperpolarization is central to and upstream of the pro-angiogenic response, we employed a chemical method, using diazoxide, to induce membrane hyperpolarization in absence of estim. This treatment caused an outflux of potassium, resulting in membrane hyperpolarization (**Figure S5**). We then assessed whether chemically-induced hyperpolarization could mimic the effects of estim on pro-vasculogenic signaling. We found that diazoxide-treated HUVECs, similarly to estim-treated HUVECs upregulated expression of the pro-vasculogenic *VEGFA* as well as *PDGFB* (**Figure S5**). Taken together, these results suggest that membrane hyperpolarization is involved in the early vasculogenic responses of the endothelial cells that we and others have observed with application of estim.

In order to test whether membrane hyperpolarization in itself is sufficient to drive increased vascularization, we used the microfluidic device model to observe changes in vasculogenesis in the presence of diazoxide. We supplemented the media cultures with diazoxide for the first 4 days to mimic the estim treatment, and fixed the devices at day 7 (**Figure S6A)**. We observed that the networks formed under chemical hyperpolarization conditions increased in the vessel density and branching nodes (**Figure S6**) in a dose-dependent manner. These results confirm that the effect of membrane hyperpolarization itself may be partially responsible for the increase in de novo vascular assembly under estim.

### Estim acts through voltage-gated potassium channels to enhance vascularization

We demonstrated that estim induces membrane hyperpolarization in endothelial cells (**Figure 4A** and **4B**) and aimed to identify the specific ion channels involved in this response. To this end, we quantified DiBAC_4_(3) fluorescence in HUVECs cultured in 2D monolayers and subjected to estim with and without exposure to various ion channel inhibitors. DiBAC_4_(3) fluorescence showed varying degrees of changes in response to estim with and without iberiotoxin (IBTX), a calcium-activated potassium channel (K_Ca_) inhibitor, tetraethylammonium (TEA) and 4-aminopyridine (4AP), both K_V_ inhibitors, quinidine (Quin.), an inward rectifier potassium channel (K_ir_) as well as voltage-gated sodium (Na_V)_ channel inhibitor, and verapamil (Ver.) as an L-type calcium channel inhibitor (**Figure 4C**). Of the inhibitors tested, only TEA and 4-AP effectively diminished the hyperpolarization effect induced by estim (**Figure 4C, Figure S4**), suggesting that K_V_ channels are crucial for membrane hyperpolarization during estim treatment. We tested for the expression of three TEA- and 4AP-responsive K_V_s (K_V_1.1, 1.3 and 1.5) and compared their expression to two channels whose expression in endothelial cells is well-established: K_ir_2.1, and TRPV4, as well as an endothelial cell marker, CD31. We found that all three of these voltage-gated channels are expressed on the surface of HUVECs (**Figure 4D**), where they would indeed facilitate the hyperpolarization effect.

Following these findings, we next investigated how inhibition of HUVEC membrane hyperpolarization via ion channel blockage affects vascular assembly and formation of perfusable networks. We employed the microfluidic devices and supplemented the HUVEC-NHDF coculture media either with vehicle control or potassium channel blockers (TEA and 4-AP) for an hour prior to start of estim and throughout the four days of estim treatment. The devices were imaged at day 7 to evaluate the changes in vascular architecture and functionality. While we observed no significant differences in the structure of the networks between the experimental conditions, we observed that functional perfusion was only detected in the estim-vessels culture in vehicle control conditions (**Figures 4E** and **4F**). Quantification of the degree of perfusion revealed a significant increase in the percentage of perfused vessels in estim control, but this effect was diminished when the devices were exposed to TEA and 4AP. This finding suggests that potassium outflux through K_V_ channels is necessary for the assembly of perfusable vasculature in response to estim.

## Discussion and conclusion

In this study, we investigated exogenous electrical stimulation as a versatile tool for improving the induction of vasculogenesis in engineered human tissues. To do this, we made observations using two modular estim devices that were designed to be cost-effective, easy to fabricate and adaptable to a variety of culture configurations. While previous studies in the field have shown that endothelial cell monolayers cultured with electrical stimulation respond by activation of pro-angiogenic pathways [10], [13], [16], [17], [18], we further built on these findings by exploring the impact of estim in a 3D *in vitro* context. Specifically, we show that endothelial cells embedded within a fibrin hydrogel together with fibroblasts assemble into a 3D vascular network which is structurally denser and more interconnected when formed under estim exposure. Additionally, we show that in a microfluidic model of vasculogenesis and anastomosis, the vessels formed in the presence of estim are significantly more perfusable.

To evaluate the translational potential of this approach, we tested the engraftment of estim-preconditioned grafts *in vivo*. Recent advances in tissue engineering have enabled the construction of many different types of tissues including heart, kidney, and liver. However, building a stable vasculature within these tissues still remains challenging. Previous studies in the field have shown that while current vascular engineering methods allow for anastomosis, the generated vessels do not persist *in vivo*. A study by Ben-Shaul and colleagues demonstrated that engineered vascular constructs require two-week preculture prior to implantation for sustained anastomosis[30]. In another approach, Zeinstra et al. demonstrated a successfully perfused vascular patch with only 4 days pre-culture, however the grafts required a complex multi-step fabrication process[31]. We show that vascular grafts pre-conditioned with estim for two days and implanted into mice are actively perfused at 14 days post-implantation while control implants are not, suggesting an improved ability of estim-exposed constructs to lumenize and anastomose with the host vessels. We further demonstrate the use of estim to vascularize ectopic liver grafts and show that we can detect active perfusion already at day 7 post-implantation in estim-pretreated grafts as well as human albumin secreted into mouse serum. This establishes a proof-of-concept with estim as an easily adaptable tool for rapid vascularization of engineered tissues.

Further studies towards engineering other tissue types will need to evaluate how the parenchymal cells respond to the estim, particularly in tissues where ion transport is tightly regulated, such as the heart or kidney. A recent report by Lu and colleagues demonstrated increase in endothelial sprouting in an model of engineered heart tissue that was cultured under estim [32]. While the model applied different DC estim parameters (increasing pulse frequency over time) than our current report (constant 1 Hz frequency throughout culture), and was not capable of achieving lumenization and perfusability, it demonstrates the promise in adaptation of our technique to other tissues. The modular nature of our system, amenable to both *in vitro* and *in vivo* contexts, renders it well equipped to investigate its utility in other tissue types.

In this context, one potential limitation of our work is that we have not fully explored the boundaries of estim approach, particularly with respect to the range of estim parameters and possible construct configurations. In this study, the electrical stimulation parameters ranges were selected based on previous reports in the field [11], [12], [13], [19], [20] and subsequently optimized for cell viability and in our 2D setup before application into the 3D context. While this ensured construct health, it leaves room for further optimization, such as systematic variation of the waveform properties and stimulation duration, to maximize vascular-specific outcomes like perfusability, maturation, or host integration. On the construct side, future studies could investigate the impact of tuning the extracellular matrix composition with both natural and synthetic polymers such as collagen, gelatin methacryloyl, polyethylyne glycol, or even decellularized organ-specific matrices, further allowing to tailor the biophysical and biochemical cues [33], [34], [35], [36]. Endothelial cell density, as well as the ratio and identity of mural cells, may also determine the extent of perfusable network formation under estim. In this study, we included fibroblasts as a supporting cell type in our constructs, which have been shown to mimic mural cell functions through secretion of pro-angiogenic factors and ECM remodeling via MMPs[37], processes that are enhanced by estim[16]. Future iterations could explore the inclusion of other mural cell types such as pericytes and smooth muscle cells, as well as investigate estim’s effects on organ-specific endothelium. By further altering the construct composition, estim may serve as a powerful addition to the current vascularization toolbox and be used not only as a stand-alone strategy but also as a complementary tool alongside established vascularization techniques to achieve optimal vascular self-assembly, maturation, and integration of engineered vasculature with host tissues.

Given the observed functional benefits of estim, we next investigated how exogenous electrical stimulation directly affects the endothelial cell electrophysiology. On a cellular level, electric fields affect the cells by movement of charged cytoplasmic molecules, modulation of membrane protein conformation, activation of voltage-sensitive transporters, protein and lipid raft distribution, electroosmosis across gap junctions, and ion transport across membranes[5], [26], [27], [38]. These processes rely on the existence of an electrical charge gradient. We found that HUVECs respond to estim by membrane hyperpolarization and demonstrated that K_V_ channels mediate this change. In our microfluidic model, inhibition of K_V_ channels, prevented estim-mediated HUVEC membrane hyperpolarization, and abrogated the observed functional improvement of the estim-formed vasculature. This suggests that membrane hyperpolarization is a key intermediate, and may point to K_V_ channels playing a specific role in endothelial biology. Additionally, we show that pharmacological membrane hyperpolarization alone leads to a pro-angiogenic endothelial gene expression profile, and enhances formation of dense and branching vascular networks.

Although K_Ca_ and K_ir_ channels are classically associated with endothelial function particularly in responses to shear stress and vasodilation where they also mediate membrane hyperpolarization [39], [40], our findings highlight the role of K_V_ channels in driving estim-mediated V_mem_ change and vascularization. Given that V_mem_ is tightly regulated via a complex arrangement of ion channels, estim-mediated V_mem_ change may also be modulated by other channels. Interestingly, the estim hyperpolarization effect was enhanced by Quinidine, an inhibitor which blocks K_ir_ channels, and a potent inhibitor of Na_V_ channels. This could be attributed to reduced compensatory influx of sodium, and highlights the complex interplay of ion fluxes when establishing V_mem_. Consistent with our report however, K_V_ channels have previously been linked to estim-dependent differentiation, suggesting their broader involvement in estim-mediated cell responses[17]. The results of our study suggest that K_V_ channels and membrane hyperpolarization may therefore be an interesting drug target in vascularization efforts and a mechanistic clue to blocking or promoting angiogenesis *in vitro*, and in clinical applications. Given that ion channel blockers are already widely used in medicine, their adaptation to focus on *de novo* vessel formation benefits should be re-evaluated particularly for regenerative medicine.

Overall, V_mem_ modulation represents a broadly conserved mechanism of tissue regulation, influencing both developmental and repair processes [4], [5], [41]. Cells with high proliferative capacity or reduced differentiation tend to be more depolarized, while terminally differentiated cells typically exhibit more hyperpolarized resting potentials[28]. Disruption of polarized gradients due to ion channel mutations has been implicated in developmental abnormalities[42]. Furthermore, hyperpolarization has been shown to suppress tumorigenesis in models of oncogene-induced neoplasia[29], [43], and recent work by Chen et al. demonstrated that K_ir_2.1-mediated fibroblast hyperpolarization promotes hair growth[44]. Together, these studies highlight the instructive role of V_mem_ across a wide range of biological contexts. Our work contributes to this growing field by highlighting the utility of exogenous estim as a tool to guide these bioelectric processes, offering possible translational potential across a wide range of regenerative applications beyond vascular biology.

Together, our work establishes exogenous electrical stimulation as an effective and broadly applicable strategy for promoting functional vascularization of engineered tissues. We show that this estim-driven effect is achieved through K_V_ channel-mediated membrane hyperpolarization. These findings reveal a powerful role of exogenous bioelectric modulation in guiding vascular assembly, and highlight the enormous potential of electrical stimulation as a broadly applicable and easily implementable tool for vascularizing engineered tissues.

## Acknowledgements

This work was supported by the by the NIH (EB033821), and the Wellcome Leap Human Organs Physiology and Engineering (HOPE) Program. S.N.B. is a Howard Hughes Medical Institute Investigator. The authors would like to acknowledge Dr. Heather Fleming, Dr. Edward Tan, Dr. Sandra March, Dr. Alan J. Grodzinsky and Dr. Ellen Roche for thoughtful discussions on experimental design and technical assistance. The authors would also like to acknowledge Dr. Susanna Elledge for editing the manuscript. The authors declare the following competing financial interest(s): S.N.B. reports compensation for consulting or board membership by Amplifyer Bio, Catalio Capital, Earli Inc., Impilo Therapeutics, Matrisome Bio, Ochre Bio, Port Therapeutics, Ropirio Therapeutics, Satellite Bio, Sunbird Bio, Vertex Pharmaceuticals, and Xilio Therapeutics. All the other authors declare no competing interests.

## Materials and Methods

### Cell cultures and preparation

Pooled donor Human Umbilical Vein Endothelial cells (Lonza, 2 different lots) were cultured in Endothelial Growth Medium – 2 (EGM-2, Lonza) and used until passage 7. RFP-Expressing Human Umbilical Vein Endothelial Cells (RFP-HUVECs, Angio-Proteomie) were cultured in EGM-2 media on flasks coated with 50 µg/mL rat tail collagen I (Corning) and used until passage 5. Neonatal Human Dermal Fibroblasts (NHDFs) were cultured in in Dulbecco’s Modified Eagle Medium with 4.5 g/L glucose (DMEM, Corning Cellgro) with 1% (v/v) penicillin-streptomycin (pen-strep, Invitrogen), and 10% (v/v) Fetal Bovine Serum (FBS, Gemini), and used before p9. Cryopreserved primary human hepatocytes (PHH; 8339, Life Technologies; ZGF, BioIVT) were aggregated with fibroblasts as described before[45] and cultured in Dulbecco’s Modified Eagle Medium with 4.5 g/L glucose (DMEM, Corning) with 10% FBS, 1% pen-strep, 15 mM HEPES (Gibco), 1% (v/v) ITS+ (human transferrin, selenous acid, and linoleic acid) premix (Corning), 7 ng/ml glucagon (Sigma), 40 ng/ml dexamethasone (Sigma) for 3 days prior to encapsulation.

### Electrode manufacturing

The gold electrodes were manufactured using electron beam metal ultra-high vacuum evaporation deposition. Soda-lime glass wafers (University Wafers) were pre-cut into desired square dimensions and placed in the electron beam deposition (EBeamTemescal FC2000). A 20 nm chromium adhesion layer was first deposited, followed by 200 nm gold layer. A 20 AW tin wire was then soldered to the surface of the electrode and PDMS (SYLGARD 184, Ellsworth) was used to passivate its surface and the electrodes were placed at 60°C overnight to allow the PDMS to cure. Each electrode was individually quality checked for electrical continuity. Before use in assembly of the estim devices, the electrodes were sterilized by autoclaving.

### Cell culture chamber assembly and seeding of 2D and 3D cultures

For 2D experiments, HUVECs and NHDFs were cultured until 90% confluency in EGM-2 media (Lonza) or in DMEM with 4.5 g/L glucose (Corning Cellgro), 1% (v/v) penicillin-streptomycin (Invitrogen), 10% FBS media respectively, washed and lifted using TypLE-express (Sigma), resuspended in phenol red free EGM-2 media (Lonza) or in the case of NHDFs, DMEM (Gibco), 1% pen-strep, 10% FBS media and seeded at density of 50k cells per well in ibidi-treated cell culture chamber slides (ibidi) and let attach overnight. The electrodes were secured to the opposite sides of each well using custom-made cut-out clips. Each well received cell-type appropriate media, additionally supplemented with vehicle control, 10 mM Tetraethylammonium chloride (TEA), 10 µM 4-amino-pyridine (4AP), 1 µM quinidine, 200 nM iberiotoxin (IBTX), 10 µM verapamil (Ver), and 1 µM or 1 nM diazoxide (for chemically-induced hyperpolarization).

The devices were placed in the incubator and the electrodes were connected to the function generator (Siglent Technologies) and checked for electrical continuity. The cell culture chambers were incubated for 1 h prior to the start of estim. Electrical stimulation was applied as 2.5 V/cm, 0.2 ms DC pulses at 1Hz and the waveform was monitored throughout the experiment using a digital oscilloscope (Siglet Technologies).

For 3D experiments, lifted HUVECs and NHDFs were resuspended in EGM-2 media and a solution of 2 million cells/mL of HUVECs, 0.25 million cells/mL NHDFs, 2.5 mg/mL fibrinogen, and 1 IU/mL of thrombin was prepared and pipetted as 20 µL droplets onto the center of each well. The devices were carefully inverted as the formed fibrin was polymerizing to avoid sedimentation of cells and incubated at 37°C for 30 min. At the end of the incubation, each well received EGM-2. The cultures were pre-incubated for 1 h prior to start of the estim.

For graft seeding, PDMS gaskets were adhered in a 0.1% BSA-coated plates. The cultures were prepared as for the 3D experiments, however were seeded in volume of 50 µL into PDMS gaskets, which were then carefully dislodged and moved into the estim devices. For experiments including hepatocyte aggregates, aggregates were added to the grafts to the final PHH concentration of 5 million cells/mL.

### Microfluidic device assembly

The molds for the microfluidic devices were made using stereolithography (Proto Labs). The molds were filled with PDMS (SYLGARD 184, Ellsworth) and cured overnight at 60°C. After cutting each individual device, they were plasma bonded to a no.1 glass coverslips (VWR) followed by annealing at 100°C for 15 min. The electrodes were inserted into their respective ports, secured with PDMS (SYLGARD 164, Ellsworth), and allowed to cure overnight at 60°C. Prior to cell seeding, the interior device chamber was plasma-activated and functionalized with 0.01% poly-l-lysine and 1% glutaraldehyde, followed by overnight wash in DI water. The next day, the devices were thoroughly dried and 0.1% BSA-coated 250 µm diameter acupuncture needles (Hwato) were inserted into the needle guides allowing for formation of top-down fabricated channels. Such assembled devices were sterilized under UV for minimum 15 min.

### Cell culture in microfluidic device

HUVECs and NHDFs were lifted using TrypLE Express (Gibco), the plates were washed with media containing DMEM, 10% FBS, and 1% pen-strep, and the cells were centrifuged at 300 x g for 5 min. A 25 mg/mL solution of fibrinogen from bovine plasma (Sigma) was prepared, and a separate solution of 1000 IU of thrombin from human plasma (Sigma). The cells were resuspended in EGM-2 media, and a solution of 3 million cells/mL of HUVECs, 0.5 million cells/ mL NHDFs, 2.5 mg/mL fibrinogen, and 1 IU/mL of thrombin was prepared and pipetted through the gel port into the bulk of the device. The devices were rotated repeatedly while the solution was crosslinking, and incubated at 37°C for 30 min in a humidified chamber. Next, EGM-2 was added to each of the media ports and further incubated at 37°C for another 15 min. The acupuncture needles were carefully removed to create the top-down fabricated channels and seeded with a solution of HUVECs at concentration of 2 million cells/mL in the incubator. The cell solution in the ports was replaced with 200 µL of fresh EGM-2 media per device, or in the case of drug treated devices EGM-2 media containing 10 mM TEA, 10 µM 4-amino-pyridine (4AP), and 1 µM or 1 nM diazoxide, and the devices were placed on a rocker and connected to the waveform generator incubated for 1 h prior to the start of electrical stimulation. Electrical stimulation was applied as 0.5 V/cm, 0.2 ms DC pulses at 1Hz and the waveform was monitored throughout the experiment using a digital oscilloscope (Siglet Technologies). The media was replaced every day of the experiment.

### Membrane potential measurements and drug treatments

HUVECS were cultured until 90% confluency in EGM-2 media (Lonza), washed and lifted using TypLE-express (Sigma), resuspended in phenol red free EGM-2 media (Lonza) and seeded at density of 50 thousand cells per well in ibidi-treated cell culture chamber slides (ibidi) and let attach overnight. On the day of the experiment, a 500 nM solution of the Bis-(1,3-Dibutylbarbituric Acid)Trimethine Oxonol (DiBAC_4_(3)) in phenol-red free EGM-2 (Lonza) was prepared. For the drug-treatment experiments, the DiBAC_4_(3)-EGM-2 was further supplemented with 10 mM Tetraethylammonium chloride (TEA), 10 uM 4-amino-pyridine (4AP), 1 µM quinidine, 200 nM iberiotoxin (IBTX, Millipore Sigma), 10 µM verapamil hydrochloride (Ver, abcam), and 1 µM or 1 nM diazoxide (for chemically-induced hyperpolarization) (Sigma), or vehicle control. Prior to the start of experiment, the gold-plated electrodes were installed in each of the wells, and DiBAC_4_(3)-containing media (with and without the added inhibitors) was added to the wells. The cells were incubated for 1 h at 37 °C prior to the start of electrical stimulation. Confocal images of DiBAC_4_(3) fluorescence were taken on the Zeiss LSM 900 microscope with 20x objective and 488 filter set at baseline, prior to the the start of estim, and at respective timepoints after continuous estim. For live imaging, the chamber devices were placed in a heated chamber and confocal images were obtained at 3, and 6 min intervals throughout the experiment over the course of 2 h of estim, with use of definite focus stabilization. The images were analyzed using Zeiss ZEN software. 10-30 individual cells were selected per field of view and the background-corrected DiBAC4(3)) mean fluorescence intensity was measured at each timepoint (F), and plotted relative to the pre-experimental, initial fluorescence (F_0_).

### Viability assay

The viability assay was conducted following treatment with estim and using Ethidium homodimer-1 (Thermo Fisher), Calcein-AM (Thermo-Fisher) and Hoechst 33342 (Thermo Fisher). A staining solution with 2 uM Calceium-AM, 1 uM EthD-1, and 8 uM Hoechst was prepared and added to cells for 15 min. The cells were imaged using Keyence BZ-X fluorescent microscope, with at least five fields of view imaged per experimental condition. QuPath 0.5.1 software was used to count the number of dead (EthD-1^+^) cells and reported as percentage of all nuclei (Hoechst^+^).

### RNA isolation and rt-qPCR

The cultures were lysed using TRIzol (Thermo Fisher) and stored at −80°C until further processing. The samples were thawed and mRNA was isolated using phenol-chloroform extraction, and the RNA-containing aqueous phase was further purified using RNA-easy min-elute cleanup kit (Qiagen), and cDNA was synthesized using RevertAid First Strand cDNA Synthesis Kit (Thermo Fisher) on a Bio-Rad CFX96 Real-Time Thermocycler, according to manufacturer’s protocol. Rt-qPCR was performed using PowerUp SYBR Green Master Mix (Thermo Fisher Scientific) on a Bio-Rad CFX384 Real-Time Thermocycler with primer sequences as listed in Table S1. Fold change gene expression was calculated using the ΔΔCT method assuming equal efficiencies, and using *GAPDH* as a housekeeping gene.

**Table S1.**
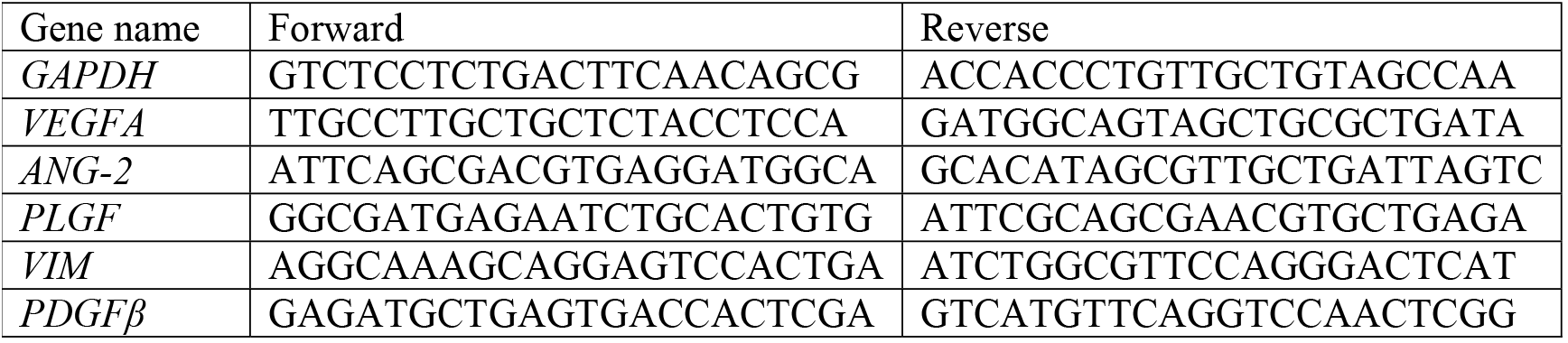
Primer sequences.

### Vessel network imaging and analysis

The cell cultures were fixed with 4% paraformaldehyde (PFA) in 1 x PBS for 30 min at room temperature (RT). All staining steps were conducted on a rocker. The samples were washed 3 times with 1 x PBS and blocked with 3% Bovine Serum Albumin (BSA) overnight at 4°C. The samples were stained with 1:500 dilution of UEA I – lectin DyLight 649 (Vector Labs) in 3% BSA overnight at 4°C, followed by 1:2000 solution of Hoechst in 1 x PBS and washed three times with 1 x PBS. Z-stack confocal images were taken using Zeiss LSM 900 and 10x and 20x objectives. Laser powers and imaging setting were kept constant within the same experimental runs. Image analysis was performed using a custom MATLAB script as described before[46], [47], any reported values are the average of the characteristic within a field of view. Images were obtained across at least two to six microfluidic devices were per condition. In the case of perfusion testing, the microfluidic devices were perfused with 0.25 mg/mL 500 kDa FITC-Dextran (Thermo Fisher) by adding the dextran into one of the media ports to create a pressure gradient and let perfuse for 30 s prior to imaging.

### Permeability Analysis

The microfluidic devices were seeded with cell-free fibrin gel, and the top-down fabricated channels were created using 250 µm acupuncture needles. The channels were seeded with HUVECs and cultured under estim for four days. The media was removed from the media ports and 10 µL of 70 kDa FITC-dextran (Thermo-Fisher) was added to only one of the ports. Fluorescent images were taken using Zeiss LSM 900 over the course of 120 seconds, at 10 second intervals. The permeability was calculated as described before[48]. Fluorescence change as a function of time was quantified in a selected region of the device close to the top-down fabricated vessel.

### Hepatocyte functional analysis

Following the estim treatment, hepatocyte viability was assessed using the RealTime-Glo™ MT Cell Viability Assay (Promega) according to the manufacturer’s protocol. The activity of CYP3A4 was measured using P450-Glo™ CYP3A4 Assay (Promega). For both assays, luminescence was read using Tecan Infinite M Plex plate reader. For albumin measurements, the culture supernatants were collected and stored at −20°C. Albumin levels in media were measured using ELISA with immobilized anti-albumin antibodies (Bethyl, cat. no. A80-129) and horseradish peroxidase– conjugated anti-human albumin antibodies (Bethyl, catalog no. A80-129P) and determined based on standard curve measurements.

### Graft implantation and mouse tissue processing

All animal procedures were conducted at Massachusetts Institute of Technology in accordance with protocols approved by the Committee on Animal Care (Institutional Animal Care and Use Committee). Female NOD *scid* gamma mice (NSG, Jackson Labs) or BALB/c nude mice (Charles River), aged five to eight weeks, were used for the implantation surgeries. Mice were anesthetized with isoflurane, and received extended-release pre-operative analgesia for post-operative pain management (extended-release buprenorphine, Ethiqa XR). Each implant was surgically placed either in the mesenteric fat pad within the intraperitoneal (IP) space or subcutaneously (SubQ). At the endpoint of the experiment (day 7 or day 14), the animals were anesthetized with isoflurane and injected retroorbitally with 70 kDa FITC-, or TexasRed-conjugated lysine fixable dextran (Thermo Fisher). The animals were euthanized, and the implanted tissues were carefully harvested. The explants were fixed in 4% paraformaldehyde (v/v) for 24 hours at 4°C. After fixation, the samples were washed with 1 x PBS, blocked with 3 % BSA solution and stained using 1:200 solution of Ulex Europaeus Agglutinin I (UEA I), DyLight 649 (Vector Labs). Confocal images of whole perfused grafts were performed using Zeiss LSM.

### Statistical analysis

All statistical tests were performed using GraphPad Prism software. One-way ANOVA test was used to compare results of experiments with more than two groups, and unpaired t-test was performed for comparison between two groups. Statistical significance was reported as p<0.05, and indicated on each graph with (*) indicating p<0.05, (**) indicating p<0.01, and (***) indicating p<0.001.

