## Supplementary Information for "Electrical stimulation directs formation of perfused vasculature in engineered tissues"

### *Supplemental Figures*

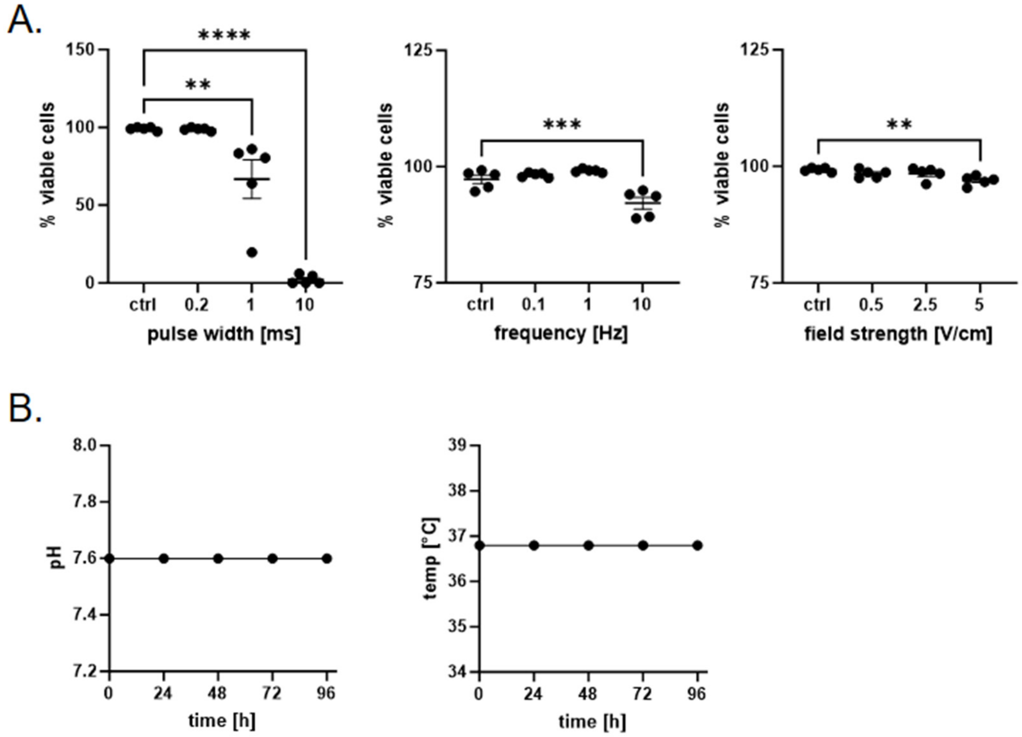


***Figure S1****. A) HUVEC viability following electrical stimulation; b) measured media pH and temperature changes over the course of 4 days of estim.*

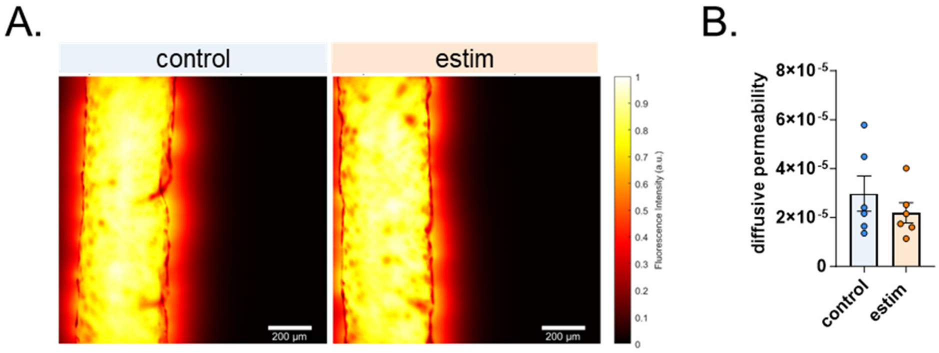


***Figure S2.*** *a) spatial heatmap of FITC-dextran diffusion out of the vessels; b) quantification of diffusive permeability in control vs estim-treated vessels.*

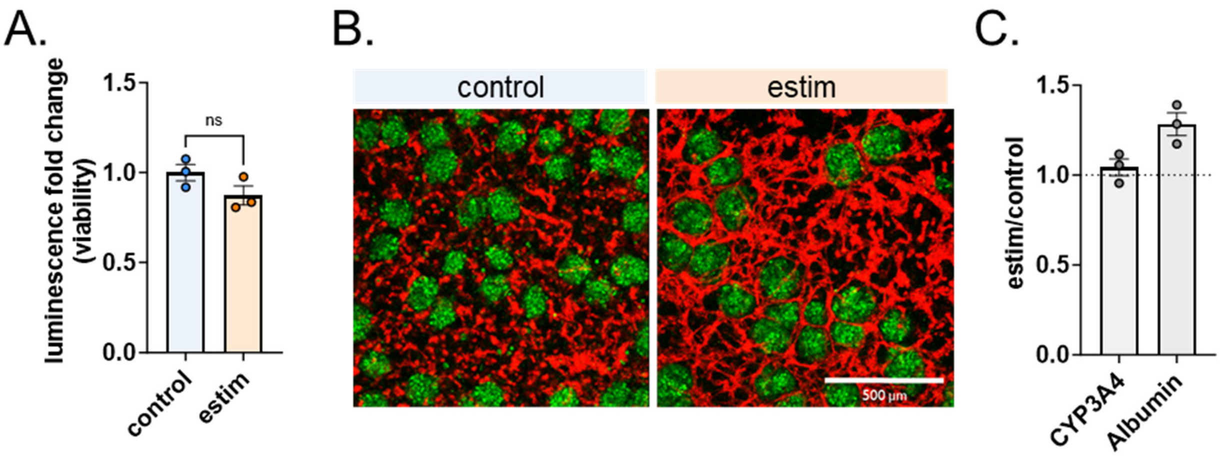


***Figure S3.*** *Hepatocytes are not negatively affected by treatment with estim. a) hepatocyte viability following treatment with estim; b) images of 3D constructs of hepatic spheroids with HUVEC; c) fold change of CYP3A4 activity and albumin secretion of human hepatocytes following estim treatment.*

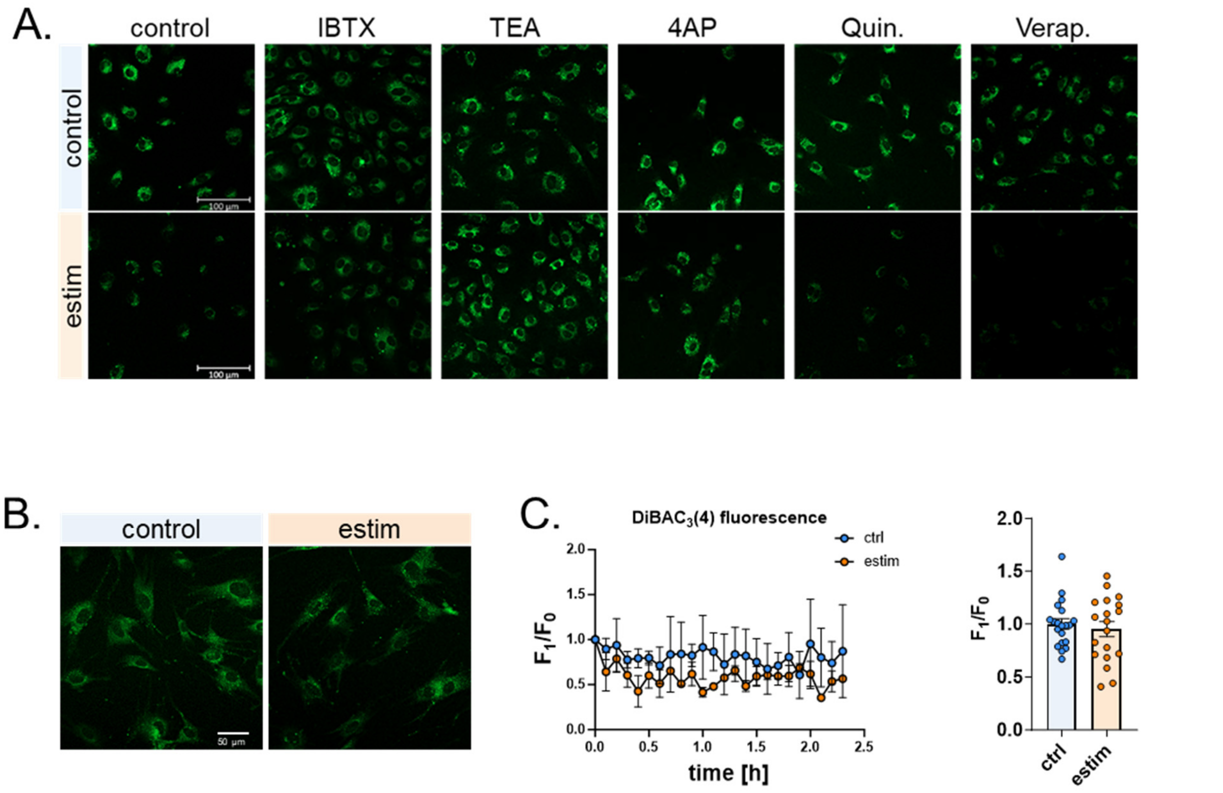


***Figure S4.*** *a) representative fluorescent images of DiBAC$_4$(3) (green) in HUVECs following estim treatment in the presence of each of the inhibitors; b) fluorescent images of DiBAC4(3) (green) in NHDFs following 2 h exposure to estim; c) quantification of changes in DiBAC$_4$(3) fluorescence in NHDFs over 2 hours of exposure to estim time and quantification of fluorescence changes in 20 individual cells at the 2 h timepoint.*

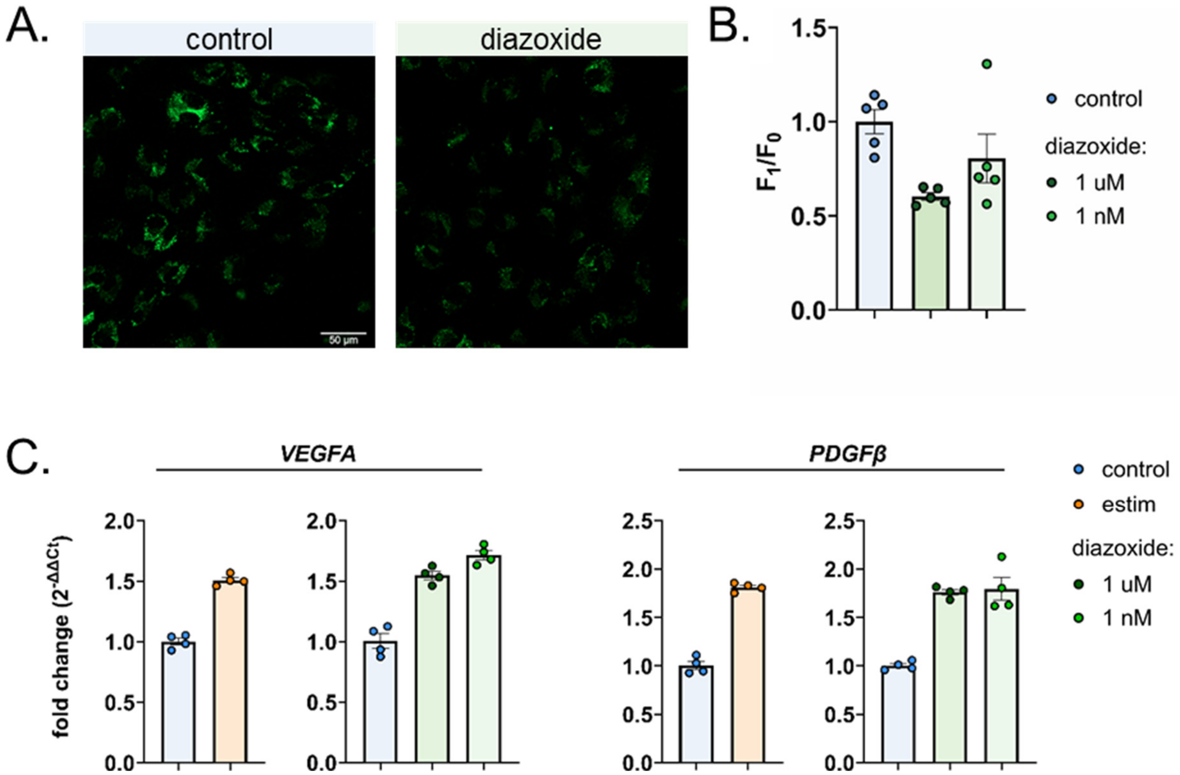


***Figure S5***. *Chemical hyperpolarization induces pro-vascular responses in endothelial cells. a) fluorescent images of DiBAC$_4$(3)-stained HUVECs (green) following diazoxide treatment; b) quantification of DiBAC$_4$(3) fluorescence changes; c) gene expression comparison changes of HUVECs treated with estim vs. diazoxide.*

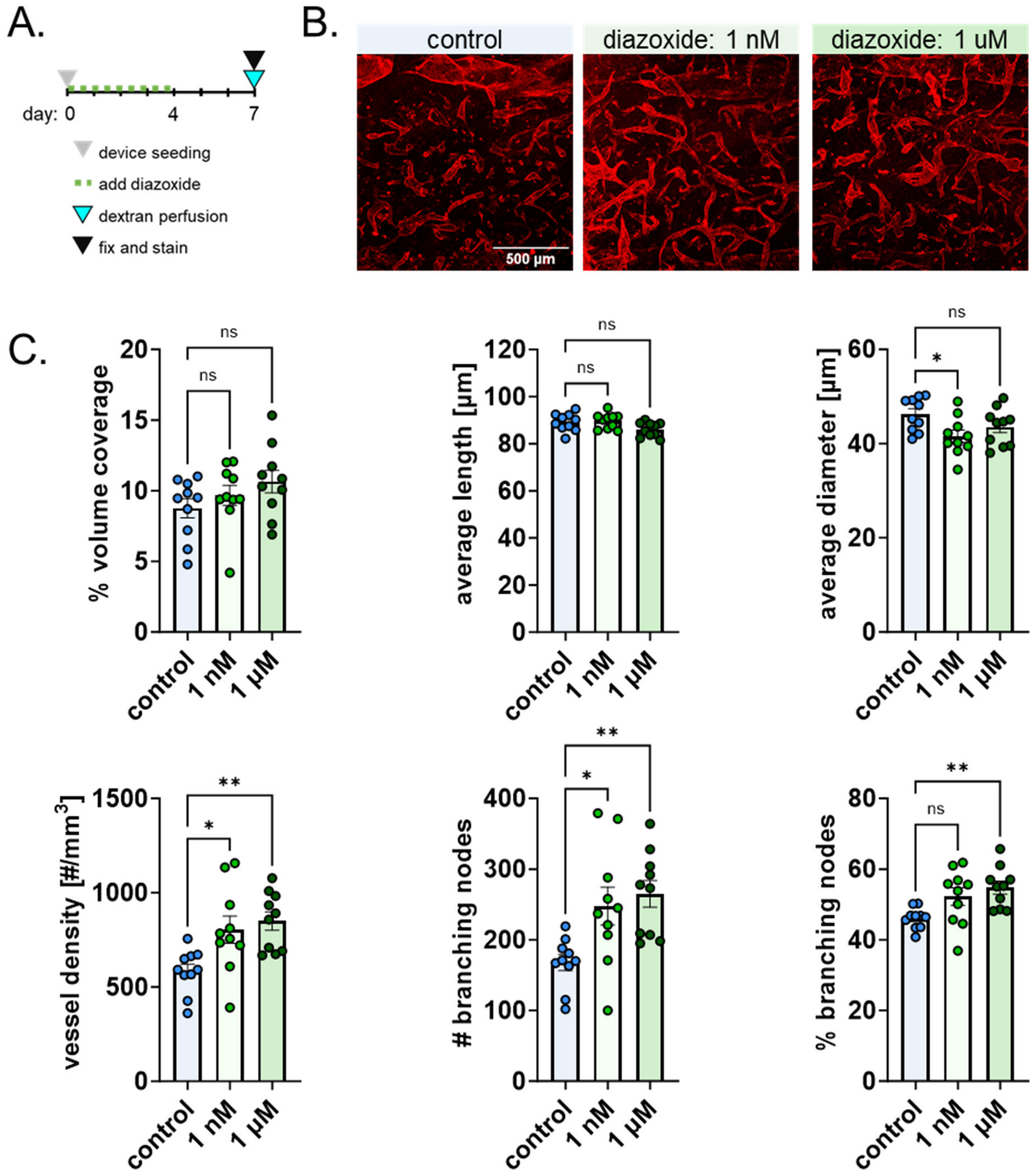


***Figure S6***. *Chemically hyperpolarized cultures form denser vasculature. a) experimental timeline; b) fluorescent images of UEA I lectin-stained HUVECs (red) in the microfluidic device following 4-days of diazoxide treatment at 7 day timepoint; c) quantification of network characteristics, ordinary one-way ANOVA, ns not significant, * p<0.05, ** p<0.01.*
